# Constrained brain volume in an efficient coding model explains the fraction of excitatory and inhibitory neurons in sensory cortices

**DOI:** 10.1101/2020.09.17.299040

**Authors:** Arish Alreja, Ilya Nemenman, Christopher Rozell

## Abstract

The number of neurons in mammalian cortex varies by multiple orders of magnitude across different species. In contrast, the ratio of excitatory to inhibitory neurons (E:I ratio) varies in a much smaller range, from 3:1 to 9:1 and remains roughly constant for different sensory areas within a species. Despite this structure being important for understanding the function of neural circuits, the reason for this consistency is not yet understood. While recent models of vision based on the efficient coding hypothesis show that increasing the number of both excitatory and inhibitory cells improves stimulus representation, the two cannot increase simultaneously due to constraints on brain volume. In this work, we implement an efficient coding model of vision under a volume (i.e., total number of neurons) constraint while varying the E:I ratio. We show that the performance of the model is optimal at biologically observed E:I ratios under several metrics. We argue that this happens due to trade-offs between the computational accuracy and the representation capacity for natural stimuli. Further, we make experimentally testable predictions that 1) the optimal E:I ratio should be higher for species with a higher sparsity in the neural activity and 2) the character of inhibitory synaptic distributions and firing rates should change depending on E:I ratio. Our findings, which are supported by our new preliminary analyses of publicly available data, provide the first quantitative and testable hypothesis based on optimal coding models for the distribution of neural types in the mammalian sensory cortices.

## Introduction

Neural circuits are responsible for a wide variety of tasks, including encoding and processing sensory information. Understanding the design principles as well as the functional computations in such circuits has been a foundational challenge of neuroscience, with potential applications to a wide variety of fields ranging from human health to artificial intelligence. However, the structural complexity and dynamic response properties of these circuits present significant challenges to uncovering their fundamental governing principles. Some of the brain’s structural properties are extremely variable across species and individuals [1], while properties such as the structure of cortical microcircuits seem to be reasonably conserved [2–5]. These conserved properties offer hope of revealing general principles of how canonical neural computations are organized.

While multiple experimental [6–13] and computational [14–16] studies have offered insights about inhibitory interneurons at different scales, their precise computational role in sensory information processing remains elusive. The relative abundance of excitatory and inhibitory neurons in primary sensory areas appears to be one of the better conserved structural properties of cortical microcircuits, and this conserved circuit structure should be an important clue for determining neural circuit function. For example, despite wide variations over several orders of magnitude in the total number of neurons across species and sensory cortical areas, morphological studies indicate that the ratios of excitatory to inhibitory neurons (E:I ratio) stay within a relatively narrower nominal range of 3:1–9:1 (i.e., inhibitory interneurons are 10% – 25% of the neural population) across species and are relatively consistent across sensory cortical areas within species (Table 1) [17–27] even when other cortical areas show variations (e.g., motor cortex [28] or medial prefrontal cortex [29]).

**Table 1:**
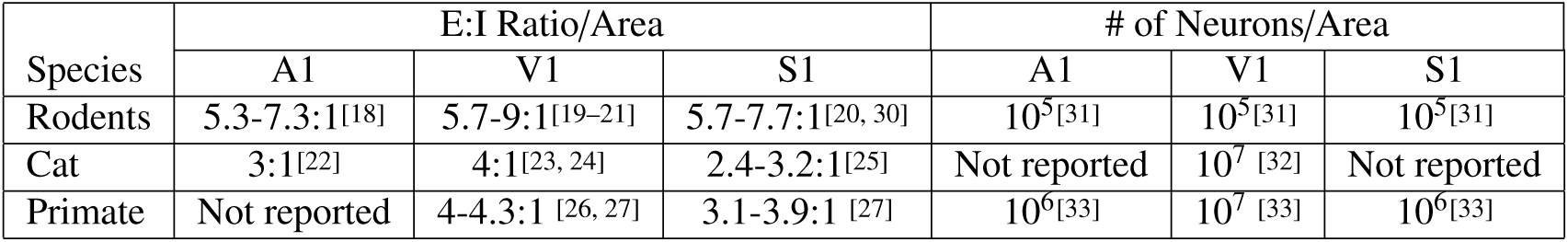
E:I ratios and # of Neurons in primary auditory (A1), visual (V1) and somatosensory (S1) cortices for different species from morphological studies.

This relative constancy of the E:I ratio must be understood within the context of sensory computations. Inhibitory interneurons in sensory cortical microcircuits have connectivity patterns contained within local circuits [2, 4], leading to inhibitory cells being generally viewed as performing a modulatory role in computation while excitatory cells code the sensory information directly [5]. For a given sensory cortical area, there are potential computational benefits to increasing the size of both the excitatory and the inhibitory subpopulations. For example, more excitatory cells may provide higher fidelity stimulus encoding, while more inhibitory cells may enable more complexity or accuracy in the computations being performed. However, volume is a critical constrained resource for cortical structures [35], and increasing one of these subpopulations in a fixed volume necessitates decreasing the other. We propose that the narrow variability of the E:I ratio can be explained as an optimal trade-off in the fidelity of the sensory representation contained in the excitatory subpopulation vs. the fidelity of the information processing mediated by the inhibitory subpopulation. Understanding this trade-off may play a critical role in determining the principles underlying the structure and function of cortical circuits.

Specifically, we propose to understand this trade-off in the context of efficient coding models [34, 36–38] under a volume constraint. In this initial study, the volume constraint is defined as the total number of neurons and does not explicitly model either volume differences by cell type or non-somatic elements such as axons and dendrites (though those extensions could be added in the future). We implement an efficient coding model known as sparse coding [34, 39], which uses recurrent circuit computations to encode a stimulus in the excitatory cell activities (denoted *a*_*j*_) using as few excitatory neurons as possible (i.e., having high population sparsity). In detail, the sparse coding model proposes encoding a stimulus (e.g., an image) *I* in terms of the sum of the activity *a*_*j*_ of excitatory neurons with receptive fields *ϕ* _*j*_, by minimizing a cost function that balances representation error (i.e., fidelity) with the sparsity of the neural population activity:

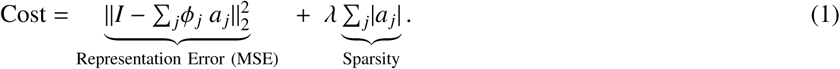

Note that the population sparsity constraint only includes excitatory cells and does not include the activity of the inhibitory cells necessary to enact the required computation (i.e., solve the optimization program). Sparse coding models have been shown to account for many observed response properties of the visual cortex [34, 39, 40] and can be implemented in biophysically plausible recurrent circuits [41, 42] with a desired sparsity level and a given E:I ratio [14, 15] (optimally approximating the ideal circuit implementation). See *Methods* for details. Recent work has shown also that increasing the population of excitatory [15, 43] and inhibitory [15] cell types in sparse coding models can improve stimulus representation in models where the size of neural populations is unrestricted.

Here we show that, for a fixed neural population size (representing a volume constraint), there exists an optimal E:I ratio where the stimulus representation, the sparseness of the sensory representation, and the metabolic efficiency of the entire network are all optimized in the model. This model-optimal E:I ratio is consistent with observed biophysical ranges and it varies based on the sparsity level of the encoding, potentially accounting for species specific variations within the observed biophysical ranges. Furthermore, higher optimal E:I ratios (at higher sparsity levels) produce inhibitory synaptic distributions that are more specific while approximately preserving the total inhibitory influence in the circuit (to retain balanced levels of excitation and inhibition). These results constitute specific and testable theoretical predictions requiring comparative neurophysiology and neuroanatomy experiments for full validation. We also perform novel analyses of experimental recordings of neural populations in area V1 for multiple species (mice, cats and monkeys), constituting the first steps in comparative analyses of population sparsity in largescale electrophysiology recordings. The results of this analysis are consistent with the model prediction of a correlation between E:I ratio and population sparsity level. Taken together, these results suggest that a combination of optimal coding models with physical constraints (e.g., volume) may provide a potential normative explanation for conserved structures observed in sensory cortical microcircuits across species.

## Results

We analyze sparse coding models optimized for a variety of E:I ratios (i.e., the ratio of the number of excitatory cells to inhibitory cells while fixing the total number of neurons) and sparsity levels (denoted by model parameter *λ*) by unsupervised training using a natural image database [34]. See *Methods* for details. The performance of these models is quantified using stimulus reconstruction error, population sparsity [45], and metabolic energy consumption [44].

For a sparse coding model trained with the sparsity constraint *λ*=0.15, we observe that the reconstruction error is minimized at the ratio of ∼6.5 : 1 (Fig. 2 (top)). The reconstruction error is a surrogate measure of the fidelity of the stimulus information preserved in the encoding. As the E:I ratio increases from 1:1, the increase in E cells leads to greater receptive field diversity in the E cell subpopulation [15, 43], allowing for better encoding of the stimulus. This increased representational capacity produces a gradual decline in the reconstruction error. As the E:I ratio increases beyond the optimum, the declining number of inhibitory interneurons results in insufficiently diverse inhibition to accurately solve the desired encoding, leading to a rapid increase in the reconstruction error. Results are independent of the size of the used database (10 images with 512 x 512 pixels each) used for training (See Fig. S2 and *Supplemental Methods*). The tolerance in calculating the optimal reconstruction error was negligible compared to the changes in the error due to varying the E:I ratio.

**Figure 1:**
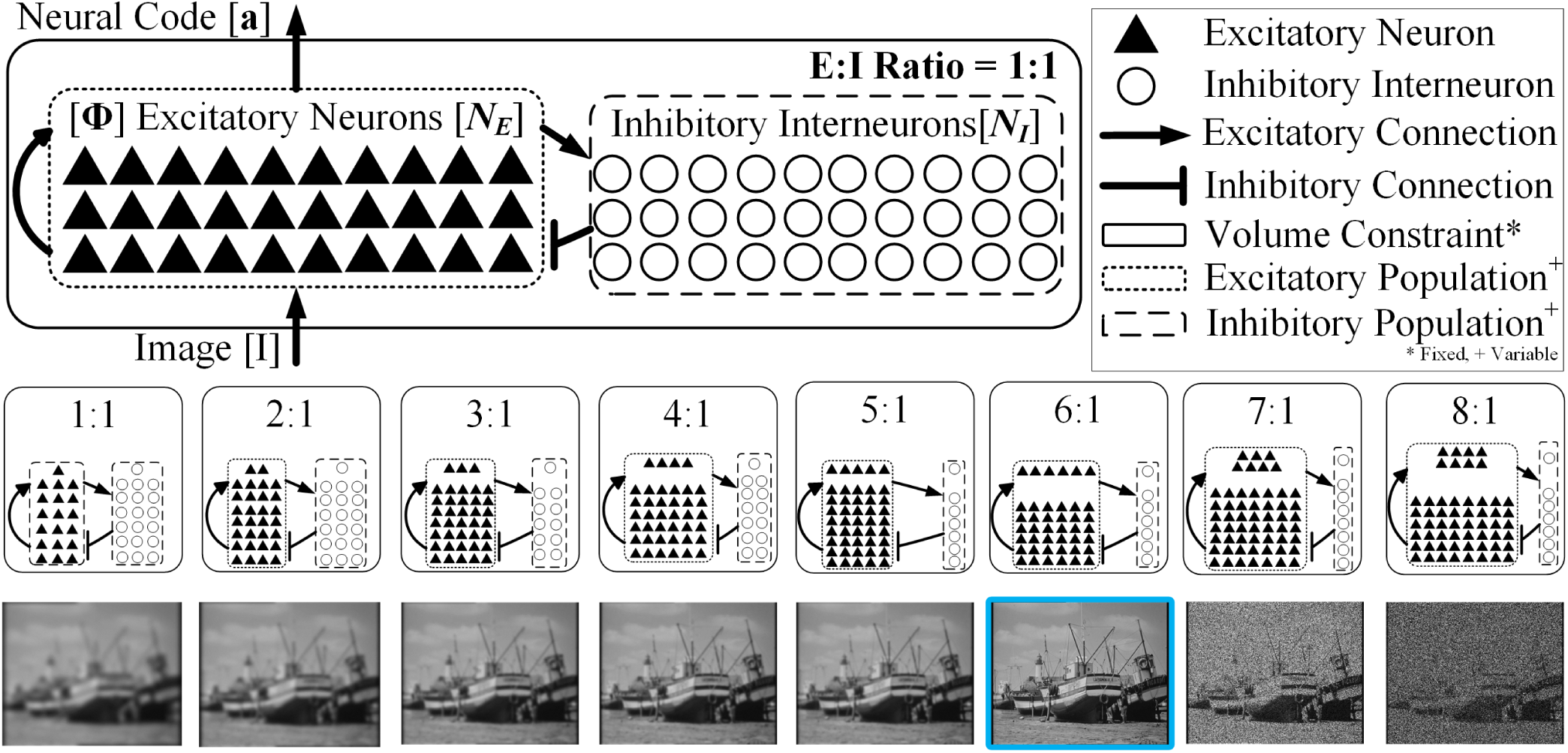
Optimal E:I ratio for coding fidelity: **(top row)** A sparse coding model is placed under a volume constraint by restricting the total number of neurons to *N*. Excitatory neurons receive recurrent as well as feed forward (stimulus) input and are responsible for coding the stimulus. Inhibitory interneurons are driven by recurrent excitatory inputs, and enable accurate computation of the neural encoding to induce sparsity in the excitatory neurons. **(middle row)** We vary the relative size of the excitatory (*N*_E_) and inhibitory (*N*_I_) subpopulations and evaluate the model at different E:I ratios under the volume constraint, *N* = *N*_E_ + *N*_I_. **(bottom row)** We show that coding fidelity is optimal (boxed image at 6:1) at a unique, biologically plausible E:I ratio for the fixed volume. We evaluate models coding 16 × 16 = 256 pixel natural image patches [34] with *N* = 1200 (≈ 5× overcomplete representation).

**Figure 2:**
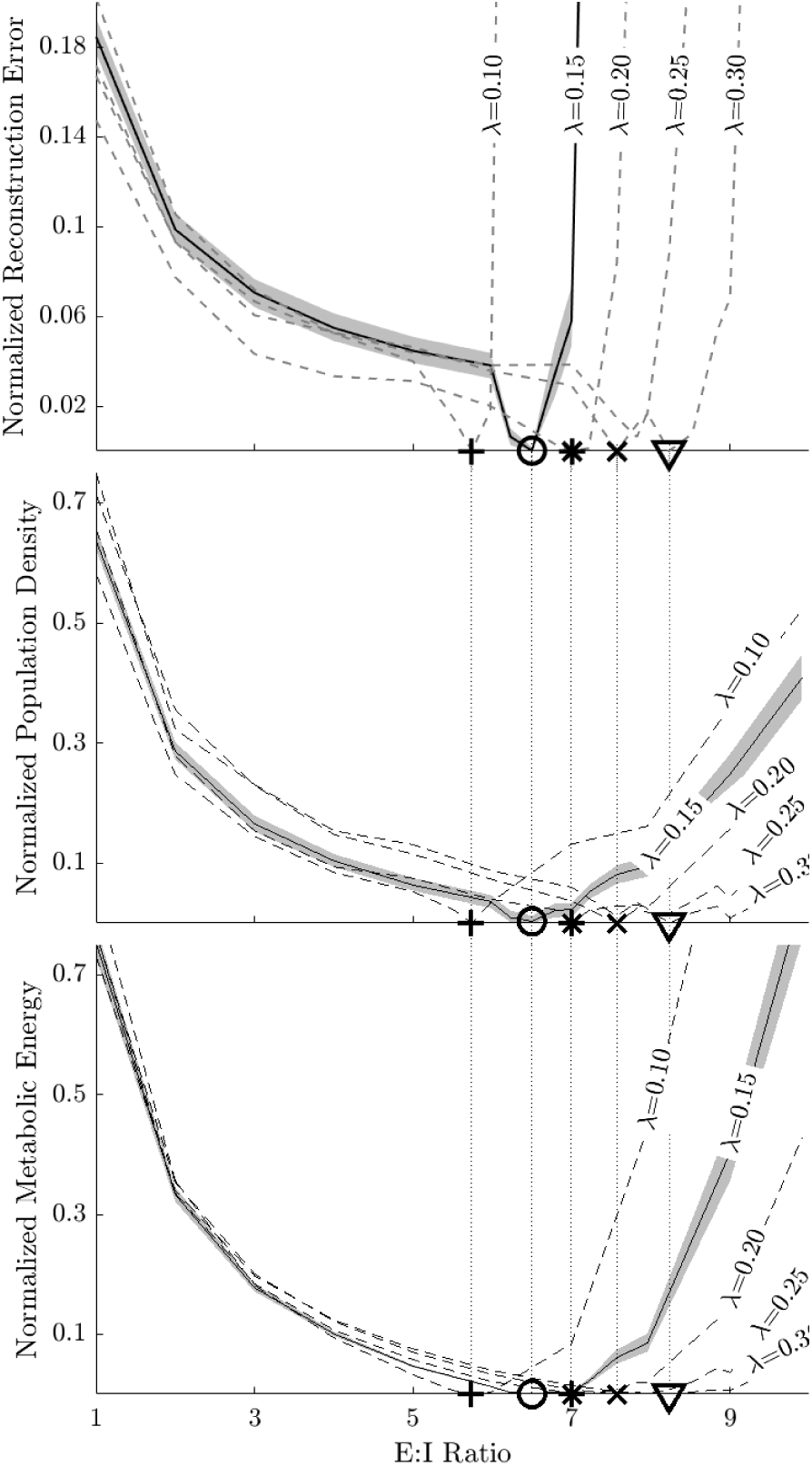
Optimal E:I ratios for multiple performance measure coincide and increase as sparsity (*λ*) increases: The performance of sparse coding models subject to a volume constraint of *N* = 1200 neurons and under different sparsity constraints (*λ* ∈ [0.10, 0.30]) and using stimuli (100 image patches, 16 x 16 pixels) drawn from a database of 10 natural 512 x 512 pixels images [34]. Performance measures are normalized per Equation 8 and standard error (depicted for *λ* = 0.15 with a shaded band) over the natural image database is estimated using a bootstrap procedure (see *Supplemental Methods*). Markers denote the optimal E:I ratio for models at each sparsity constraint for each performance measure. Optimal E:I ratios for different performance measures are essentially identical as illustrated by vertical lines connecting markers across the 3 plots, and increases in model sparsity (*λ*) correspond to increases the optimal E:I ratio for each performance measure.**(top)** The coding fidelity for a sparse coding models with different sparsity constraints quantified by the normalized reconstruction error. The coding performance is optimized at an E:I ratio of approximately 6.5:1 (in a biophysically plausible range), with values above (below) that number suffering from lack of diversity in the inhibitory (excitatory) cell population. **(middle)** Population Activity Density (1 - Population Sparsity) for a sparse coding model (see *Methods*) is minimized at nearly the same specific optimal E:I ratio as with coding fidelity. **(bottom)** Lastly, a metabolic energy consumption measure [44] (see *Methods*) reveals minimal metabolic energy consumption at nearly the same specific E:I ratio as with coding fidelity and population density.

Efficient coding models seek a parsimonious representation of sensory inputs in the excitatory neural activity in addition to an accurate encoding. To quantify this parsimony, we plot the density of activity of excitatory neurons in the sparse coding model (Fig. 2 (middle)), as measured by population density, an additive inverse of the commonly used modified Treves-Rolls (TR) metric [45] that quantifies population sparsity (see *Methods*). Notably, the population activity density is minimized (i.e., population sparsity is maximized) at approximately the same E:I ratio that optimizes reconstruction fidelity. At low E:I ratios, the stimulus representation is not rich enough to admit a sparse representation of natural scene statistics with available receptive fields of excitatory cells. With high E:I ratios, the available inhibition is insufficient to achieve sparse population activity in the excitatory cells.

A common rationale for the efficient coding hypothesis (including sparse coding models) is that efficient codes may reduce the metabolic cost of the neural activity [46–49]. While decreasing the mean firing rate of excitatory neurons would decrease the metabolic cost of producing action potentials in those cells, it is not clear which network architecture minimizes the total metabolic energy consumption when accounting for the cost of supporting the non-sparse activity of the inhibitory interneurons [9, 50, 51]. We quantify and plot (Fig. 2 (bottom)) the total metabolic energy cost of the network (see *Methods*). Once again, the optimal E:I ratio achieving minimal energy consumption for different sparsity constraints is approximately the same E:I ratio that optimizes reconstruction fidelity and excitatory population sparsity.

When varying the sparsity constraint across a wide range *λ* = [0.1, 0.3], we observe that all three performance measures (reconstruction error, population sparsity, metabolic energy consumption) demonstrate an optimal E:I ratio that is consistent across metrics. This indicates that there is a clear optimal E:I ratio for a given sparsity level that is robust to the choice of optimality criteria. Crucially, we observe that increasing the model sparsity (*λ*) leads to a higher optimal E:I ratio in all three metrics (Fig. 2).

While networks optimized for different sparsity levels have different optimal E:I cell type ratios, it is unclear if either the synaptic distribution (a structural measure) or the total amount of inhibitory activity (a functional measure) change as well. To understand potential structural changes, we first examined the structural nature of the inhibitory interactions in the recurrent network at different sparsity levels (*λ*) and optimal E:I ratios. We observe that there are systematic changes in the distribution of weights for Inhibitory →Excitatory connections (Fig. 3 (middle row - left)) as *λ* changes. In particular, lower sparsity levels (*λ*) corresponding to lower optimal E:I ratios result in inhibitory synapse distributions that have heavier tails and higher kurtosis (Fig. 3 (middle row - right)). Therefore, at lower E:I ratios when there are relatively more inhibitory interneurons in the circuit, the individual interneurons have more targeted projections to deliver inhibition more selectively to shape excitatory activity (Fig. 3 (top row), see also Fig. S3 (left) and (right)).

**Figure 3:**
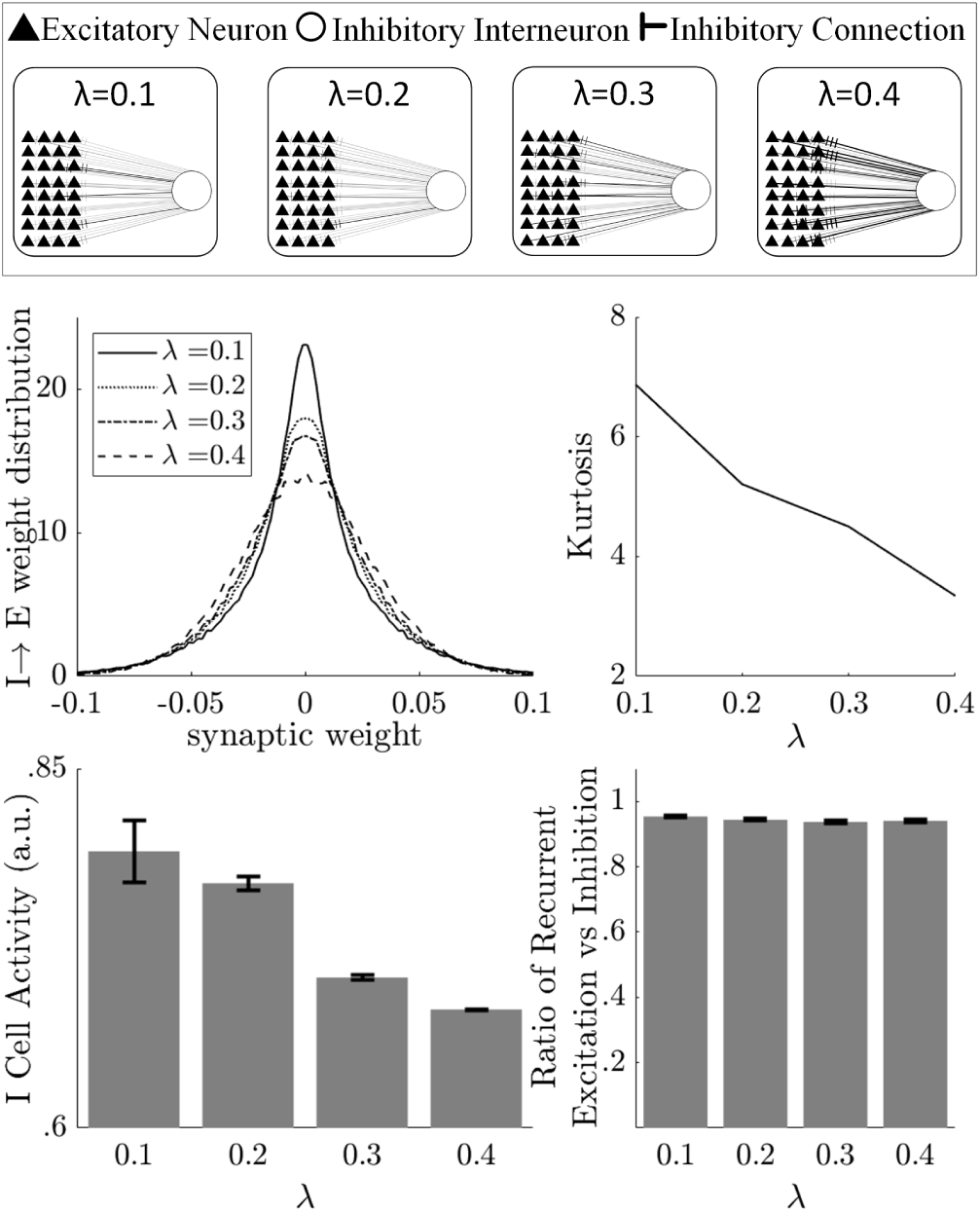
Structure and function of model inhibition change with sparsity: **(top row)** An illustration visualizing the impact of the changing weight distributions on an inhibitory interneuron. **(middle row) (left)** Estimated probability density functions for the inhibitory to excitatory connection weights in the optimal computational models at different sparsity levels reveal an increasing fraction of inhibitory synapses are stronger as sparsity increases. **(middle row) (right)** Estimated kurtosis vs sparsity quantifies the changes visible in the distributions, demonstrating that inhibition is more targeted and less global at lower sparsity levels with smaller E:I ratios. **(bottom row) (left)** With increasing sparsity (corresponding to higher optimal E:I ratios), the inhibitory subpopulation’s mean activity level declines and becomes less diverse (exhibiting a lower standard deviation). **(bottom row) (right)** Despite the changes in inhibitory structure and function due to changes in sparsity level (and optimal E:I cell type ratio), the changes to inhibitory synaptic distributions and firing rates counteract each other so that the total inhibitory influence in the network remains constant and the circuit maintains balance between the recurrent excitatory and inhibitory activity.

Functionally, the total amount of inhibitory influence in a circuit is a combination of the spiking activity in the inhibitory interneurons and the total strengths of the synapses from inhibitory to excitatory neurons. We next examined the inhibitory activity in the recurrent network at different sparsity levels (*λ*) and optimal E:I ratios. We observe that lower *λ* corresponding to lower optimal E:I ratios result in higher average activity levels per cell (with higher standard deviations) across the relatively larger inhibitory subpopulation (Fig. 3 (bottom row - left)). Despite significant changes in the synaptic structure and firing rates of inhibitory interneurons as *λ* (and the E:I cell type ratio) changes, the total amount of inhibitory influence in the network does not change substantially (Fig. 3 (bottom row - right), S4). Specifically, as *λ* increases, the reduction in inhibitory subpopulation size and firing rates is offset by the broader tuning of the inhibitory synapses so that the balance between total excitation and inhibition in the network remains relatively constant in a stable regime (Fig. S3 (middle)).

While theoretical modeling often assumes that the sparsity level of an efficient coding model is an unknown parameter that can be fit to data, the analysis above predicts that optimal efficient coding networks should have E:I ratios correlated with population sparsity (Fig. 4). Unfortunately, despite sporadic characterizations of population sparsity reported in the literature (with different data types and analysis methods), we lack a comparative analysis of population sparsity across species. In new analyses of recent publicly available datasets comprised of large-scale V1 electrophysiology recordings, we evaluated the sparsity in population activity in mice [52], monkeys [53, 54] and cats [55] studies featuring natural visual stimuli (movies, images). The similarly low sparsity levels observed in monkeys and cats (both having E:I = 4:1) as well as their contrast with higher sparsity levels in mice (E:I = 5.7-9:1) are consistent with the predictions of the efficient coding model in this study. Specifically, using a hierarchical bootstrap procedure [56] (See *Methods, Supplemental Methods* and Fig. S1) to compare population sparsity for different species, we observed (Fig. 4 (bottom row) that mice have much higher population sparsity (lower density) than monkeys and cats when viewing natural movies (*p*_*bootstrap*_ < 10^−8^). Similarly, mice exhibit higher population sparsity than monkeys (*p*_*bootstrap*_ = 0.01966) in response to natural images.

**Figure 4:**
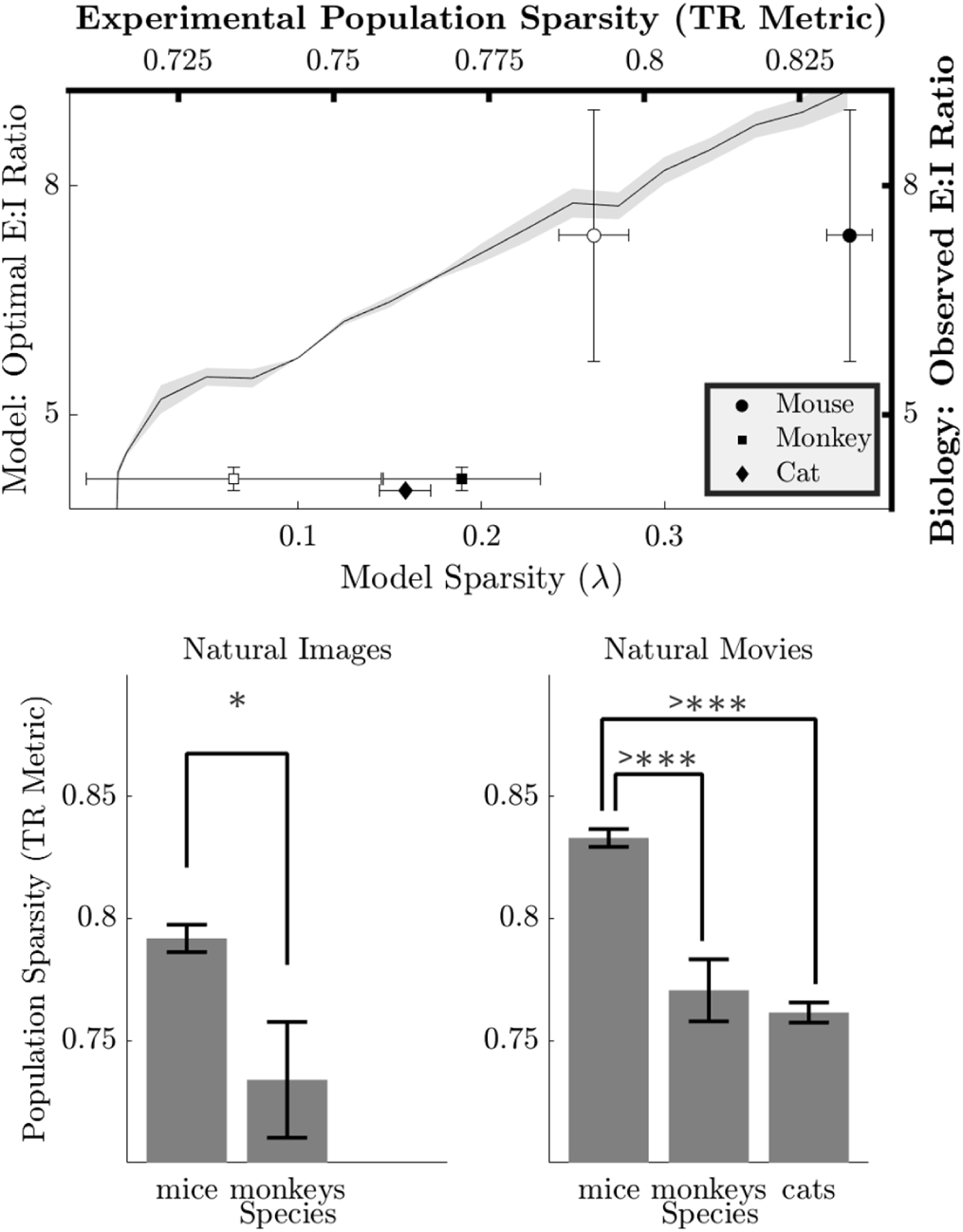
Model predictions vs experimental data (Top Row): **(Left Y and Bottom X axes)** Optimal E:I ratio based on normalized reconstruction error (See Fig. S5 for other performance measures) as a function of model sparsity constraint *λ* is depicted by the solid line (mean) with variability (± standard error) denoted by the shaded band. **(Right Y and Top X axes)** The population sparsity (TR) measure computed for electrophysiology data from experimental studies in mice [52], monkeys [53, 54] and cats [55] is shown (mean (markers) ± standard error (horizontal error bars)) as a function of observed E:I ratio ranges in biology (vertical error bars). Unfilled markers represent natural images and black filled markers represent natural movies. **Interspecies comparisons (Bottom Row)** Statistical significance of hypotheses based on model prediction (i.e., higher E:I ratio in biology corresponds to higher population sparsity) examined via inter-species population sparsity comparisons with all available data using hierarchical boostrapping. **(left)** For natural images, the mice (E:I = 5.7-9:1) exhibit higher population sparsity compared to monkeys (E:I = 4-4.3:1), *p*_*bootstrap*_ = 0.01966. **(right)** For natural movies, mice (E:I = 5.7-9:1) exhibit higher population sparsity than both monkeys (E:I = 4-4.3:1) and cats (E:I = 4:1), *p*_*boostrap*_ < 10^−8^ for both, which is significant after accounting for multiple comparisons.

## Discussion

Using a sparse coding model for early vision and a volume constraint, we showed that the quality and efficiency of stimulus encoding is optimal at E:I ratios consistent with the narrow range observed in biological neuroanatomy. Increasing the E:I ratio improves the representational capacity of the E cell subpopulation through the potential for greater receptive field diversity [15, 43], but at the expense of reducing the ability of the I cells to produce accurate circuit computations to implement the encoding rule. Decreasing the E:I ratio has an opposite effect, increasing the I cells available to improve computational accuracy for the encoding rule at the expense of the representational capacity of the E cell subpopulation, whose receptive field diversity shrinks, diminishing its ability to represent rich sensory statistics.

This model makes several predictions that are testable with comparative electrophysiology experiments. The primary result of this study predicts that the optimal E:I ratio is directly correlated with population sparsity, such that sparser population activity in a species will correspond to a higher E:I cell type ratio (Fig. 4). In secondary results, this model also predicts that species with higher sparsity levels will have inhibitory interneuron subpopulations with both lower average firing rates that are more concentrated around the mean and higher kurtosis of the synaptic distribution than species with lower sparsity levels. These predictions are notable because it is rare for computational theories to make specific and measurable predictions about the relationship between functional and morphological properties of neural systems.

The result that networks with a higher level of population sparsity in the excitatory subpopulation are optimized with fewer inhibitory neurons (i.e., higher E:I ratio) may appear counter-intuitive given the apparent need for increased inhibition to achieve higher sparsity. However, a closer look at the specific structure in the inhibitory synaptic distribution (see *Methods* and *Supplementary Information*) provides some insight into this result. Models having higher population sparsity learn to represent natural stimuli differently from models at lower population sparsity. Specifically, in models with higher population sparsity, the smaller inhibitory subpopulation contains cells that have relatively lower firing rates and global synaptic connections, indicating inhibition that is more broadly tuned and less selective than in models with lower population sparsity. This model prediction is consistent with the contrast observed in experimental results from cats (E:I = 4:1) [57] and mice (E:I=5.7-9:1) [58]. In contrast, models at lower population sparsity have inhibitory interneurons with relatively higher firing rates and synapses that are targeted to specific excitatory sub-populations (Fig. 3). We note that while we discuss inhibitory interneurons generally here, we have not attempted to correspond the inhibitory components of the model to a specific genetic subtype of inhibitory interneuron. Future experimental tests of the predictions from this model can and should address the empirical question of which inhibitory interneuron subtypes are the best fit to the inhibitory influences of this model.

To perform a preliminary evaluation of this model prediction with data that is currently available, we analyzed population sparsity in area V1 of mice [52], monkeys [53, 54] and cats [55] using publicly available electrophysiology data sets. We found that the population sparsity trends revealed by this analysis agree with the broad predictions made by the model. Specifically, for a given stimulus type, species with higher E:I ratios demonstrated higher population sparsity levels. To our knowledge this is the first comparative analysis of population sparsity across species, providing valuable insight for future computational and theoretical work beyond the specific predictions of this model.

Despite this apparent agreement between experimental data sets and model predictions, the predicted correlation between optimal E:I ratio and population sparsity is challenging to thoroughly evaluate empirically because the literature currently lacks the necessary reports to provide a substantive comparative analysis of population sparsity between species. The large scale populations recordings necessary to evaluate sparsity have only become possible relatively recently, and comparability of existing studies is often hampered by differences such as recording methodology, experimental conditions (e.g. type and quantity of anaesthesia administered), brain area, number of subjects, number and type of neurons, stimuli, and analysis parameters (e.g. window size substantially influences sparsity measures). The data we analyzed come from experiments whose design was not aimed at facilitating comparisons like those made in this study, and experiments that control for these sources of variability may allow for more robust evaluation of our (and future) model predictions. As an example, population recordings analyzed in this study featured lightly anesthetized mice [52] compared to heavily anesthetized and paralyzed monkeys [53, 54] and cats [55]. Since anesthesia is known to depress neural activity [59–61], we anticipate population sparsity for monkeys/cats is elevated. This bias would make it more difficult to observe the significant differences in sparsity level reported in this study, so it is unlikely to be a major confound in our analysis. However, further studies that explore population level activity in different sensory areas or under different experimental conditions may support/refute whether our model predictions apply more generally.

To illustrate the challenges with making comparative meta-analyses from data that was not collected for that purpose, we note that in addition to the data supporting the model predictions above, we have also encountered a limited number of contrasting exceptions that have known confounds that highlight the subtleties in such comparative analyses. For example, one study [62] captures V1 responses to natural stimuli in ferret and reports population sparsity (TR = 0.42) much lower than cats and monkeys despite a higher E:I ratio of 5:1 [63]. However, this study self-identifies a critical methodological issue that likely resulted in overestimated firing rates due to the use of multi-unit signals instead of isolated single units to compute sparseness, deflating the estimated population sparsity. For another example, [64] captures population sparsity in mouse V1 using spike trains estimated from calcium imaging and reports a lower population sparsity (TR = 0.45-0.55) than a recent calcium imaging study [65] (TR = 0.81), as well results from analysis of electrophysiology data from mice presented in this paper. Closer examination of this inconsistency reveals that [65] features specific targeting of excitatory neurons only while [64] does not employ cell-specific targeting, which can deflate population sparsity estimates due to the elevated firing rates of inhibitory interneurons [9, 50, 51]. The confounding effects present in these two conflicting examples from the literature illustrate a number of important methodological issues to be carefully addressed in future experimental work that aims to perform a conclusive comparative analysis.

The results of this study represent an early step toward understanding the connection between optimal coding rules and the diversity sensory cortical structure in mammals. We expect that additional verifiable predictions will be possible when more relevant biological details are introduced into the models. For example, our analysis does not make distinctions between different kinds of inhibitory interneurons and future work may consider their relative contributions when evaluating the trade-off between computational accuracy and representational capacity. Similarly, modeling thalamic input into inhibitory cells may offer greater insight into the role of inhibition beyond modulating computation performed by the excitatory sub-population.

Finally, we note that the shape of the performance curves (Fig. 2) are asymmetric, with performance degrading very quickly at E:I ratios higher than the optima. While normative models can never ensure they are capturing all constraints that drive evolutionary or developmental goals for a system, this asymmetry indicates that the constraints considered here are more robust to decreasing E:I ratios rather than increasing E:I ratios. This prediction is consistent with the (limited) currently available morphological data (Table 1) that shows the distribution of E:I ratios across species is asymmetric and skewed to smaller values around the mode. Additional morphological studies on animal models not listed in Table 1 may provide additional support or refutation of this prediction. More broadly, we expect that close interplay between computational and experimental studies will further advance our ability to merge functional and physical constraints to better understand the relationship between the information processing in the brain and its structure.

## Methods

### Sparse coding model of visual computation

Among neural coding models instantiating the efficient coding hypothesis, we concentrate on the sparse coding model [34] that aims to minimize the number of simultaneously active neurons for each stimulus. This model is sufficient to explain the emergence of classical and nonclassical response properties in V1 [34, 42, 66] and is consistent with recent electrophysiological experiments [67–69]. Furthermore, the sparse coding model can be implemented in recurrent network architectures with varying degrees of biophysical plausibility [38, 42, 70–72], including distinct inhibitory interneuron populations [14, 15].

Specifically, in the sparse coding model, a set of neurons encodes an image intensity field *I*(*x, y*) through the vector of activities *a* = [*a*_1_, *a*_2_, …] (i.e., firing rates) by minimizing the so called *energy function*:

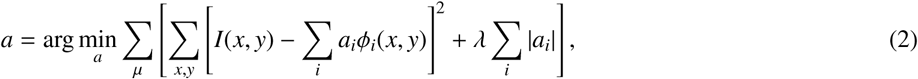

where the activity of each neuron *a*_*i*_ is associated with a stimulus feature *ϕ*_*i*_(*x, y*) (similar to a receptive field), and µ = 1 … *M* sums over all images in a training set. This energy function uses the scalar parameter *λ* ∈ [3.78 × 10^−4^, 0.4] to balance the preservation of stimulus information (measured by the mean-squared reconstruction error in the first term) with the efficiency of the representation (measured by the sum of the activity magnitudes in the second term). We choose the *L*_1_ norm for quantifying the efficiency of the representation since it is known to promote sparsity and is (analytically and computationally) tractable. Higher values of *λ* encourage more sparsity and lower values prioritize the fidelity of the stimulus encoding. As has been shown in the past, optimizing the feature set *ϕ*_*i*_(*x, y*) for this coding rule using a corpus of natural images will produce a set a features that resemble the measured receptive fields in primary visual cortex [34, 42].

### Dynamical System implementation of the sparse coding model

To encode a specified image, we consider a recurrent dynamical circuit model [70] that provably solves the optimization in Eq. [2] [73, 74] (including alternative sparsity penalties [75]) in non-spiking or spiking [71, 76, 77] network architectures. Specifically, the system dynamics for this encoding model are:

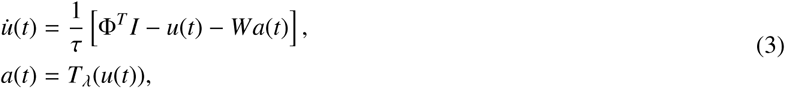

where *I* is the vectorized version of the stimulus, Φ is a matrix with a vectorized version of the dictionary element *ϕ*_*i*_(*x, y*) in the *i*^*th*^ column, the vector *u* contains internal state variables (e.g., membrane potentials), the vector *a* contains external activations (e.g., spike rates) of excitatory neurons that represent the stimulus, the matrix *W* governs the connectivity between the neurons (requiring inhibitory interneurons for implementation), and *T*_*λ*_() is a pointwise nonlinear activation function (i.e., a soft thresholding function).

When the recurrent influences in the network are governed by *W* = *G* − *D* = Φ^*T*^ Φ − *D*, where *G* is a Grammian matrix and *D* is the diagonal identity matrix, then the network above is guaranteed to converge to the solution of the sparse coding objective function above [70]. In this case, the required connectivities between the excitatory cells (the principal cells encoding the stimulus) must be mediated by a combination of direct excitatory synapses (negative elements of *G*) and a local population of inhibitory interneurons (positive elements of *G*). Deviations from this network structure may result in more efficient implementations (e. g., requiring fewer inhibitory neurons), but will have the consquence of only approximately solving the desired coding objective.

We seek to form a circuit model that approximates the ideal dynamical system above as closely as possible under a fixed size for the inhibitory interneuron population implementing *G*. To reflect the disynaptic connections onto an inhibitory population and back to the excitatory population, consider the factorization of this connectivity matrix using the singular value decomposition (SVD): *G* = *U*Σ*V*^*T*^. If we consider only the positive entries in this representation as in [14], each column of *V* contains the synaptic weights of the connections onto a single inhibitory cell, the corresponding element in the diagonal matrix Σ represents a dendritic gain term, and the corresponding column of *U* represents the synaptic weights from that inhibitory cell back onto the population of excitatory principle cells. Following previous work [14], we can use the truncated SVD to find the closest approximation (in terms of the Frobenius norm) to *G* with a specified rank, which corresponds to specifying the size of the inhibitory population.

Experimental and computational studies have reported that depending upon factors such as location, timing and magnitude, PSPs arriving at the dendritic tree can produce sub-linear, supra-linear and linear gain at the soma [78, 79]. Interpreting Σ as a gain term enables us to incorporate the biologically realistic notion of dendritic gain arising from multiple projections from an inhibitory interneuron to an excitatory neuron, into an otherwise abstract circuit model limited to representing a single projection. Under this interpretation, we estimate the activity of inhibitory interneurons as *b* = *Va*.

### Volume Constraint Implementation

For this study, we represent 16×16 pixel image patches using *N* = *N*_I_ + *N*_E_ = 1200 total neurons to correspond to a fixed volume constraint (implicitly assuming approximately constant volume per neuron). For each E:I ratio tested, we trained a dictionary using natural images [34] for a dictionary optimized for *N*_*E*_ excitatory cells. After training the dictionary, we implemented the dynamical system described above with the best approximation to the ideal circuit dynamics using *N*_*I*_ inhibitory cells.

In addition to evaluating the model at different E:I ratios, we also trained and evaluated models under different sparsity constraints (*λ*). For a given sparsity constraint (*λ*) and E:I ratio, we evaluate the network over an image patch database [39] using three different performance measures.

### Performance measures

The first performance measure quantifies the coding fidelity of the model for the reconstruction 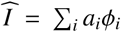 of an image *I* encoded by the model. The stimulus reconstruction error is formulated as:

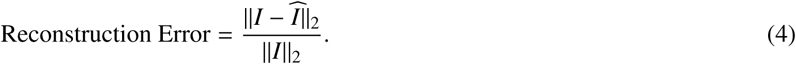

The second performance measure is population sparsity using the modified Treves Rolls (TR) metric [45]. TR scores lie between 0 and 1, with 1 being the highest sparsity. We computed model sparsity using excitatory neuron firing rates (*a*_*i*_, *i* = 1……*N*_*E*_). Existing literature on experimental evidence for sparse activity in the cortex [50] indicates that typically a small inhibitory interneuron sub-population (*a*_*i*_, *i* = *N*_*E*+1_……*N*) is far more active than excitatory neurons owing to its role in modulating activity of the entire circuit. Thus sparsity is not expected to be a feature of this sub-population, and these neurons are not included in the TR metric:

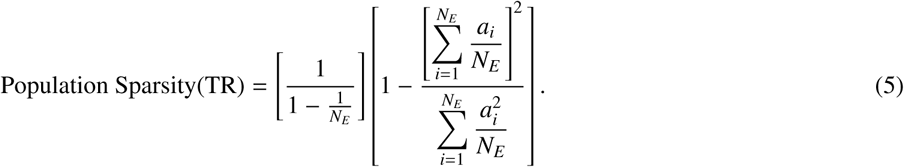

We define Population Density (or Population Activity Density) as

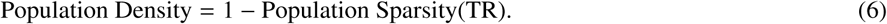

The TR metric is sensitive to bin sizes used to evaluate spike trains and smaller bin sizes lead to higher estimates of sparsity. This consideration does not affect the analysis of model activity (*a*) which is interpreted as a fixed firing rate. However, the inherent variability of spike trains in experimental data means that the choice of bin size does affect population sparsity computation. In this study, a bin size of 100ms is used for natural images, natural movies and spontaneous activity. Analysis for natural images is bound to a 100ms bin size due to a 106ms trial duration constraint in monkey experimental data [53]. A direct comparison between population sparsity of the model and experimental data is not practical given the sensitivity of the TR metric to scaling, since the dynamic ranges for the model coefficients and firing rates of neurons are very different.

The third performance measure is an estimate of the metabolic energy consumption in volume constrained sparse coding models. We compute this measure using metabolic energy consumption models for rodents and primates [44, 80], which are grounded in physiological and anatomical studies. The models estimate the metabolic energy consumption (ATP molecules/gm-minute) for cortical gray matter by aggregating estimates for the granular processes involved in its functioning. The processes include pumping out Na^+^ entering during signaling, glutamatergic signaling, glutamate recycling, post-synaptic actions of glutamate and pre-synaptic Ca^2+^ fluxes and glial cell activity. While the energy consumption associated with inhibition is thought to be somewhat less than excitation [44], we approximate the energy consumption of spiking activity as being equal in all neuron types due to the relatively smaller prevalance of inhibitory neurons and synapses in the population [44]. We have not included energy consumption due to glial cells due to their relatively small fraction of energy usage [81] and lack of a central role in the current modeling study.

For our study, we compute the metabolic energy consumption of volume constrained sparse coding models using the rodent metabolic energy consumption model, which has two main components. The first component represents the energy expended to maintain resting potentials (3.42 × 10^8^ ATP molecules/s-neuron), and the second represents energy spent to sustain action potentials at a given rate (7.1 × 10^8^ ATP molecules/neuron-spike × firing rate (Hz)). These estimates are used to compare the performance of a model at different E:I ratios, and they are only weakly affected by whether the rodent or the primate metabolic energy consumption is used:

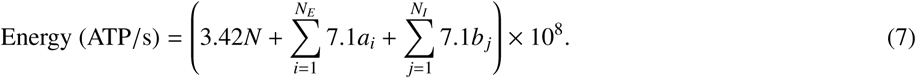

### Normalization of Performance Measures

Models with different sparsity constraints (*λ*) produce deviations against different baselines for reconstruction error, population sparsity/density and metabolic energy consumption. To compare different models, a common baseline is required. We implement normalization for each of the measures above in the form of a relative increase as a percentage of the minimal value observed across all E:I ratios evaluated for a given model. This normalization is described as

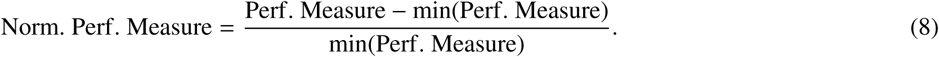

While the normalization makes visualization easier, it does not change the qualitative results.

### Inter-Species Comparisons of Experimental Population Sparsity

We computed the Population Sparsity (TR metric) for electrophysiology data sets for monkeys [53, 54], mice [52] and cats [55] that includes natural images and natural movies as stimuli types. Neural recordings from each study can be viewed as multilevel data sets, with differences in numbers of subjects, trials and neurons across them that can be represented as a hierarchy. For each trial in each data set, we compued a population sparsity value. To test model predictions that higher optimal E:I ratios correspond to greater population sparsity against experimental data from different species, we implemented a hierarchical bootstrap procedure that is more conservative in controlling for Type-I errors with multi-level data sets than traditional paired tests [56].For each species and stimulus type, we run the bootstrap 10,000 times, generating estimates of average population sparsity. We used the resulting distributions to test the hypotheses framed by model predictions. The hierarchical organization for the bootstrap procedure for each species and stimulus type is described in detail in *Supplemental Methods* and Fig. S1.

## Acknowledgements

For cat electrophysiology, neural data were recorded by Tim Blanche in the laboratory of Nicholas Swindale, University of British Columbia, and downloaded from the NSF-funded CRCNS Data Sharing website [55]. For primate electrophysiology, neural data in response to natural images were collected in the laboratory of Adam Kohn at the Albert Einstein College of Medicine and downloaded from the CRCNS web site [53]. Lastly, we appreciate feedback from Bilal Haider, Josh Siegle and Adam Kohn on the early drafts of this manuscript.

This work was partially supported by NIH R01EY019965 and NSF CCF-1350954 to C.R and NIH EB022872, NS099375, NS084844 and NSF BCS1822677 to I.N.

## Supplementary Information

### Supplementary Methods

#### Estimation of statistical errors in the analysis

We evaluate variance and bias in reconstruction error, population sparsity and metabolic energy over the image patch database of 10 natural images from which image patches are sampled for the sparsity constraint *λ*=0.15 at each E:I ratio (1:1-10:1).

To estimate the variance, we randomly select 1 out of *N* = 10 images (with replacement) and we select 10 16×16 pixel image patches from this image. Repeating this process 10 times, we gather 100 16×16 pixel image patches. We perform inference in the model and calculate the mean reconstruction error, population sparsity and metabolic energy consumption are computed using the model corresponding to each E:I ratio. This constitutes one run. We collect and aggregate statistics from 100 runs. The standard deviation of the means computed for each of the runs is the standard error for a given performance measure for a given model.

Estimation of the bias is similar, however, instead of choosing patches from *N* = 10 images, we use *N*^*^ images, where *N*^*^ = *αN, α* < 1. Mean reconstruction error, population sparsity and metabolic energy consumption are computed for each E:I ratio. This constitutes a single run. For a given *α*, we perform 100 runs. We repeat this process for each *α* ∈ {0.5, 0.6, 0.7, 0.8, 0.9, 1.0} (*α* = 1.0 being the variance estimate mentioned above), which amounts to a total of 600 runs. The average (over 100 runs) optimal E:I ratio for *λ* = 0.15 at different values of *α* is then examined to explore if the image patch database size induces any bias in the observed optimal E:I ratio.

#### Estimation of the ratio of recurrent excitation vs recurrent inhibition during stimulus representation

To better understand the effects of changes in the size of inhibitory subpopulation (i.e., different E:I ratios), we examine the relationship between recurrent excitation and recurrent inhibition received by active units in response to stimulus image patches. We consider the following decomposition of the low rank approximation (*G*) of the recurrent connectivity matrix utilized in an earlier study [1]

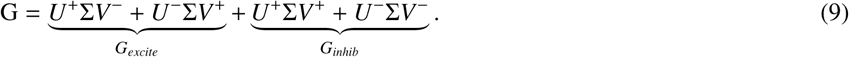

The recurrent excitation and inhibition received by active nodes is computed as

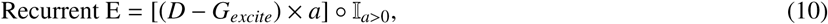

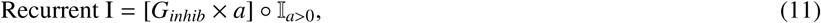

where 𝕀 is the standard indicator function taking the value 1 if the argument is true and 0 otherwise. We compute a ratio between recurrent excitation and recurrent inhibition received by active neurons as

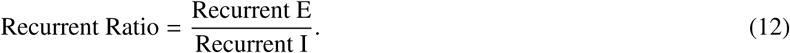

The Recurrent Ratio is computed for each anatomical E:I ratio for models with different sparsity constraints (*λ*). Here we evaluate the recurrent E/I balance specifically in response to stimulus, and recognize that the network is more stable when recurrent inhibition is greater than recurrent excitation.

**Figure S1:**
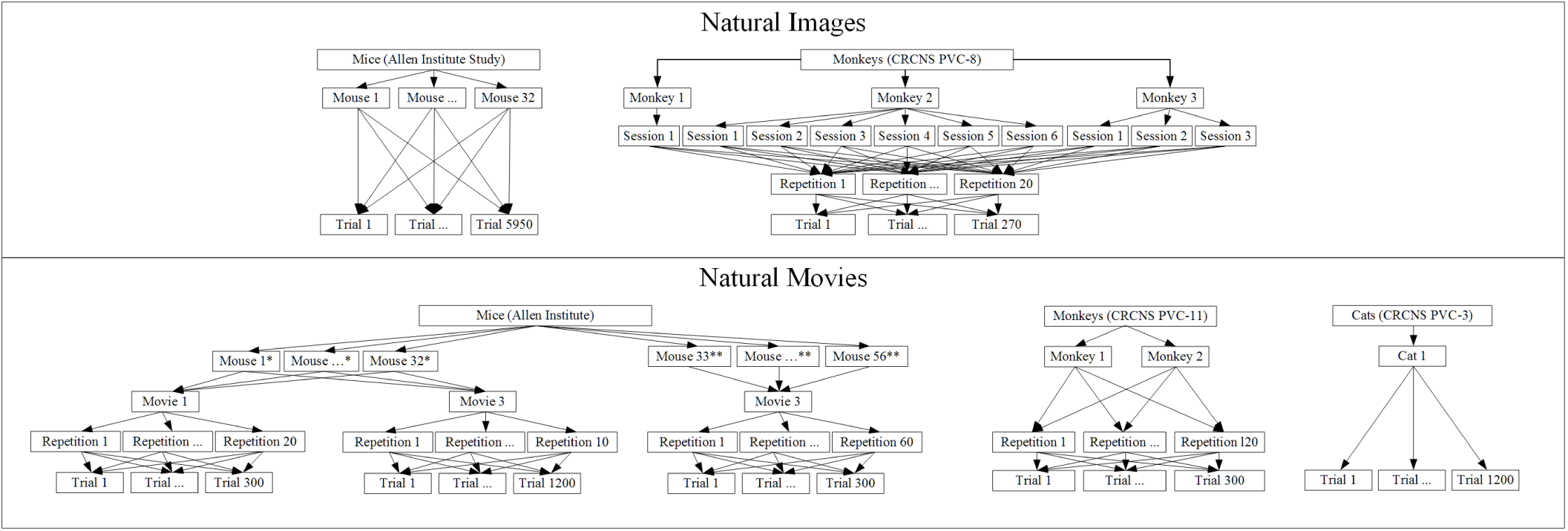
The hierarchy of multi-level experimental data sets used to draw comparisons between population sparsity of different species is represented in each figure for different stimulus types. **(top panel)** For natural images, the available data sets allow us to draw a comparison between 32 mice (E:I = 5.7-9:1) [3] with 44-244 V1 neurons and 3 monkeys (E:I = 4-4.3:1) [4] with 16-76 V1 neurons. The monkey data [4] features multiple recording sessions per subject (1,6,3 sessions for monkeys 1,2,3) and 20 repetitions of all stimuli images in a block structure. **(bottom panel)** For natural movies, the available data sets allow us to draw a comparison between 56 mice [3] with 44-244 V1 neurons, 2 monkeys [5] with 69-104 V1 neurons and 1 cat [6] (E:I = 4:1) with 10 V1 neurons. The mouse subjects come from 2 different experiments (* denotes the Brain Observatory Experiment with 32 subjects, and ** denotes the Functional Connectivity Experiment with 24 subjects) where the key differences include the number of different natural movies shown and how often each movie is repeated.

#### Hierarchical bootstrap: Hypothesis testing model predictions against multi-level experimental data

The hierarchical bootstrap procedure is built around sampling with replacement at different levels of a hierarchy in a multi-level dataset at each bootstrap run to estimate averages. The procedure is described in detail in [2].

Our measure of interest in this study is the scalar population sparsity measure (TR metric), which means that the number of recorded neurons don’t feature in our hierarchy. As an example specific to its usage in this study, we describe the case of the hierarchical bootstrap for comparing average population sparsity between mice and monkeys for natural image stimuli from multi-level data sets visualized in Figure S1 (top panel), to test the hypothesis/model prediction that mice should exhibit higher population sparsity than monkeys. For computational efficiency, population sparsity is pre-computed for each dataset before the hierarchical bootstrap procedure.

For mice, we first sample subjects (first level of hierarchy) with replacement from the 32 mice with stimulus responses to natural images in the Allen Institute data set [3]. Next, for each sampled mouse, we sample trials (second level of hierarchy) with replacement from the total number of trials ‘T’ (T=5950 in the example). Finally, we average the population sparsity across all the sampled trials to obtain an average population sparsity for natural images in mice. This process represents a single bootstrap run. A slightly different hierarchy, shown in Figure S1 (top panel) is constructed for monkeys, where the first level represents different recording sessions (with different numbers of neurons) for a single monkey. The same hierarchical bootstrap procedure is repeated. We collect average population sparsity estimates for mice and monkeys for a total of 10,000 bootstrap runs for natural images as well as other stimulus types.

For hypothesis testing related to the example above, we treat the 10,000 average population sparsity values for mice and monkeys as a 2 dimensional distribution. We use this joint distribution to evaluate the model hypothesis/prediction that population sparsity in mice (E:I = 5.7-9:1) should be higher than monkeys (E:I = 4-4.3:1). With mice on the x-axis and monkeys on the y-axis, we compute the volume of the distribution where *x* > *y* (i.e. the volume of the distribution below the line *y* = *x*). If the volume 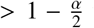 then mouse pop sparsity > monkey pop sparsity at level *α*, (*α* = 0.05 in our analysis). In the event of multiple comparisons (e.g. natural movie stimulus), Bonferroni (or other) corrections can be applied the same way as traditional hypothesis testing. The volume of the distribution opposing the hypothesis (above *y* = *x* in our example) is the *p* value for the test. It is referred to as *p*_*bootstrap*_ to disambiguate it from *p* values emanating from traditional hypothesis testing.

### Supplementary Results

#### Bias in statistical analysis

We estimated bias in all three performance measures using a bootstrap procedure (See *Supplementary Methods*) which samples from a subset of the natural image database [7] for models trained with a sparsity constraint of *λ* = 0.15. Fig. S2 indicates stability of the mean optimal E:I ratio over 100 runs sampling from differently sized subsets (denoted by *α*) of the natural image databases, suggesting negligible bias.

**Figure S2:**
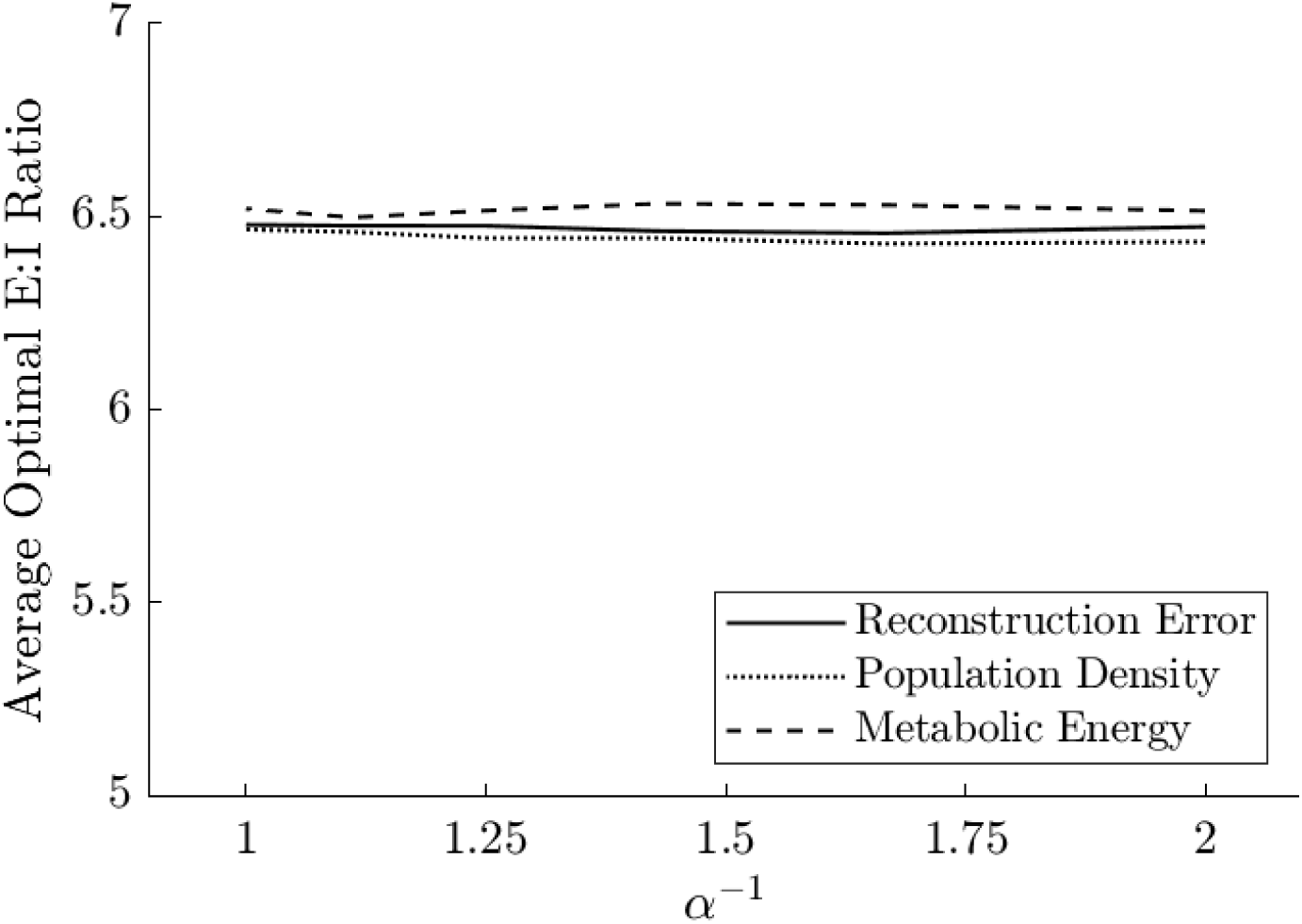
Estimation of the bias involves choosing 100 patches from *N*^*^ = *αN, α* < 1 of the *N* = 10 natural images. Mean reconstruction error, population sparsity and metabolic energy consumption are computed for each E:I ratio. This constitutes a single run. For a given *α*, we perform 100 runs. This process is repeated for each *α* ∈ {0.5, 0.6, 0.7, 0.8, 0.9, 1.0}, which amounts to a total of 600 runs. Average optimal E:I for *λ* = 0.15 and different values of *α* are shown for reconstruction error, population density and metabolic energy consumption. The relative constancy of average optimal E:I ratio over the bias runs for different values of *α* indicates that any possible bias in estimating the optimal E:I ratio is negligible for the explored sample sizes.

#### Structure of Recurrent Inhibition

We examined the static structure of inhibition of the model at the optimal E:I ratio for different sparsity levels. We observed that inhibitory strength, represented by the Singular Values Σ, interpreted as implementing dendritic gain (see *Methods*) is distributed less evenly across the inhibitory sub-population as sparsity (*λ*) increases (Fig. S3(left)), even as the total inhibitory strength/dendritic gain across the inhibitory sub-population remains relatively unchanged across different sparsity levels (Fig. S3(middle)). Next, we use a metric called the stable rank [8] which is defined as

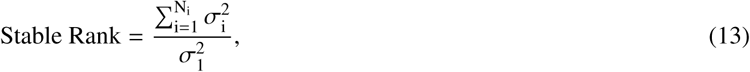

where σ_*i*_ is the *i*^*th*^ singular value of the SVD of the recurrent matrix. The stable rank is relatively robust to smaller singular values. In the context of our interpretation of Σ as the dendritic gain, the stable rank can serve as an additional measure of unevenness (lower stable rank implies greater unevenness). The value of the stable rank decreases as sparsity (*λ*) increases (Fig. S3(right)) adding support for the preceding result that indicates that unevenness in inhibitory strength of the model at the optimal E:I ratio increases as sparsity increases. Together, these results suggest that while the total amount of inhibition supported by the structure is relatively unchanged for models at optimal E:I ratio at different sparsity levels, it is distributed less evenly in the inhibitory sub-population.

**Figure S3:**
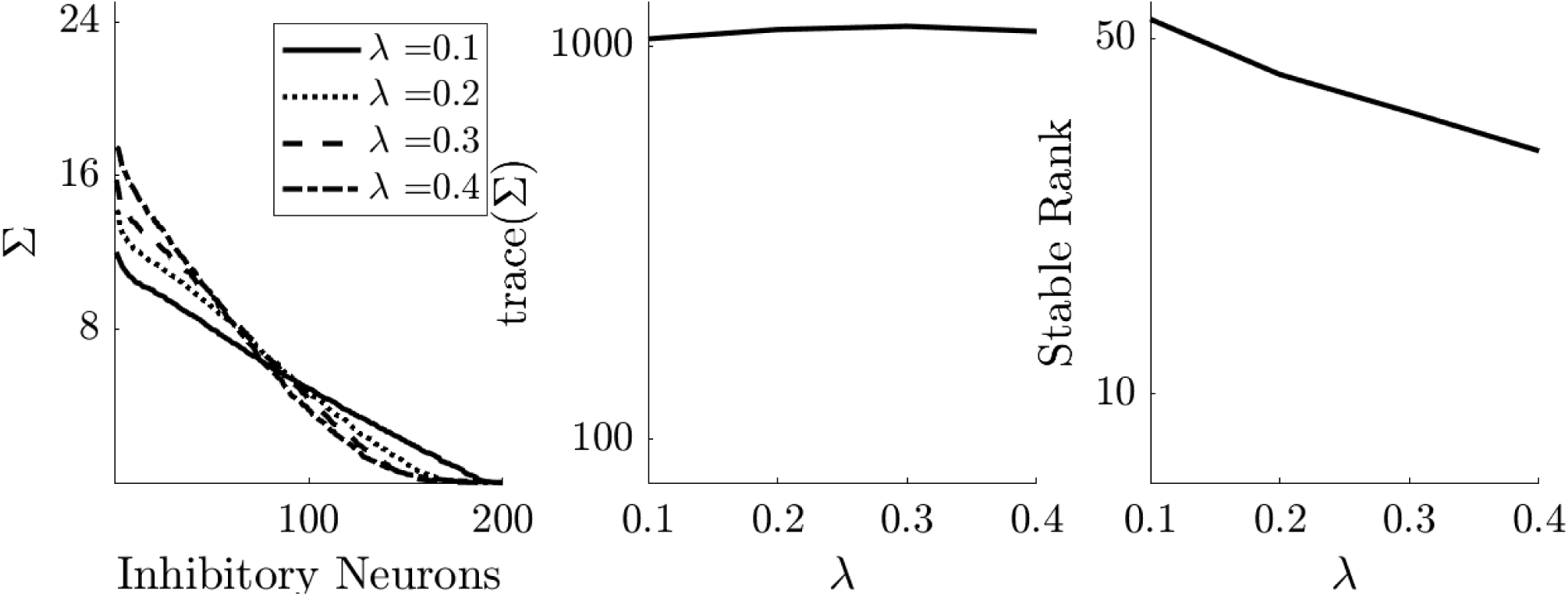
**(left)** Inhibitory strength Σ interpreted as being implemented via dendritic gain (see *Methods*) vs Inhibitory Interneurons (Components) for different sparsity levels (*λ*). **(middle)** Total amount of inhibitory strength/dendritic gain across the inhibitory sub-population, relatively unchanged for models at the optimal E:I ratio for different sparsity constraints (*λ*). **(right)** The trend of increasing unevenness in inhibitory strength as sparsity (*λ*) increases, depicted by the first two plots is also reflected by a decrease in the stable rank measure detailed in [8].

#### Inhibitory sub-population activity profiles at different sparsity levels

**Figure S4:**
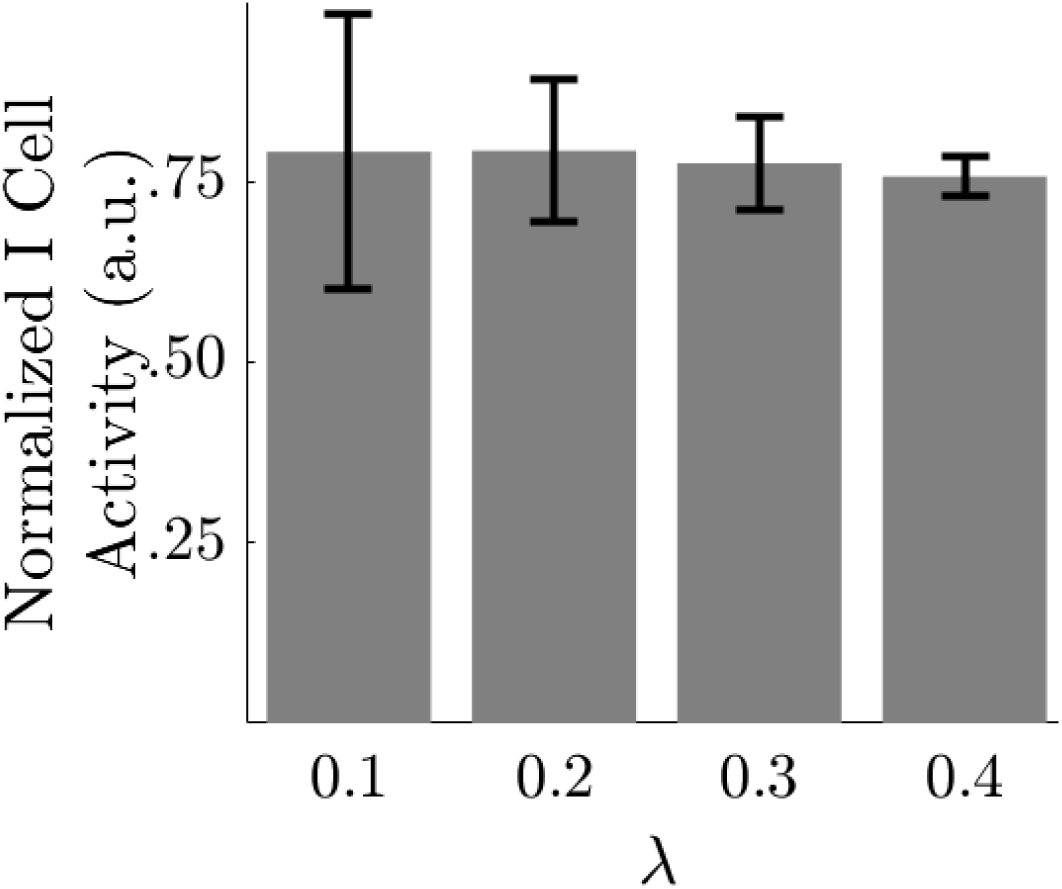
A normalized version of the (bottom row)(left) plot in Fig. 3 shows I cell activity when normalized against the total (E+I cell) activity of the model. The normalized I cell activity is relatively unchanged across optimal models at different sparsity levels, while the diversity of responses (error bars) to different natural image stimuli in the inhibitory sub-population shows (like the un-normalized plot) that I cell responses are more specifically tuned to stimuli at lower sparsity levels/lower optimal E:I ratio and become broadly tuned and less diverse as model sparsity/optimal E:I ratio increases.

#### Model performance (all performance measures) vs biology

**Figure S5:**
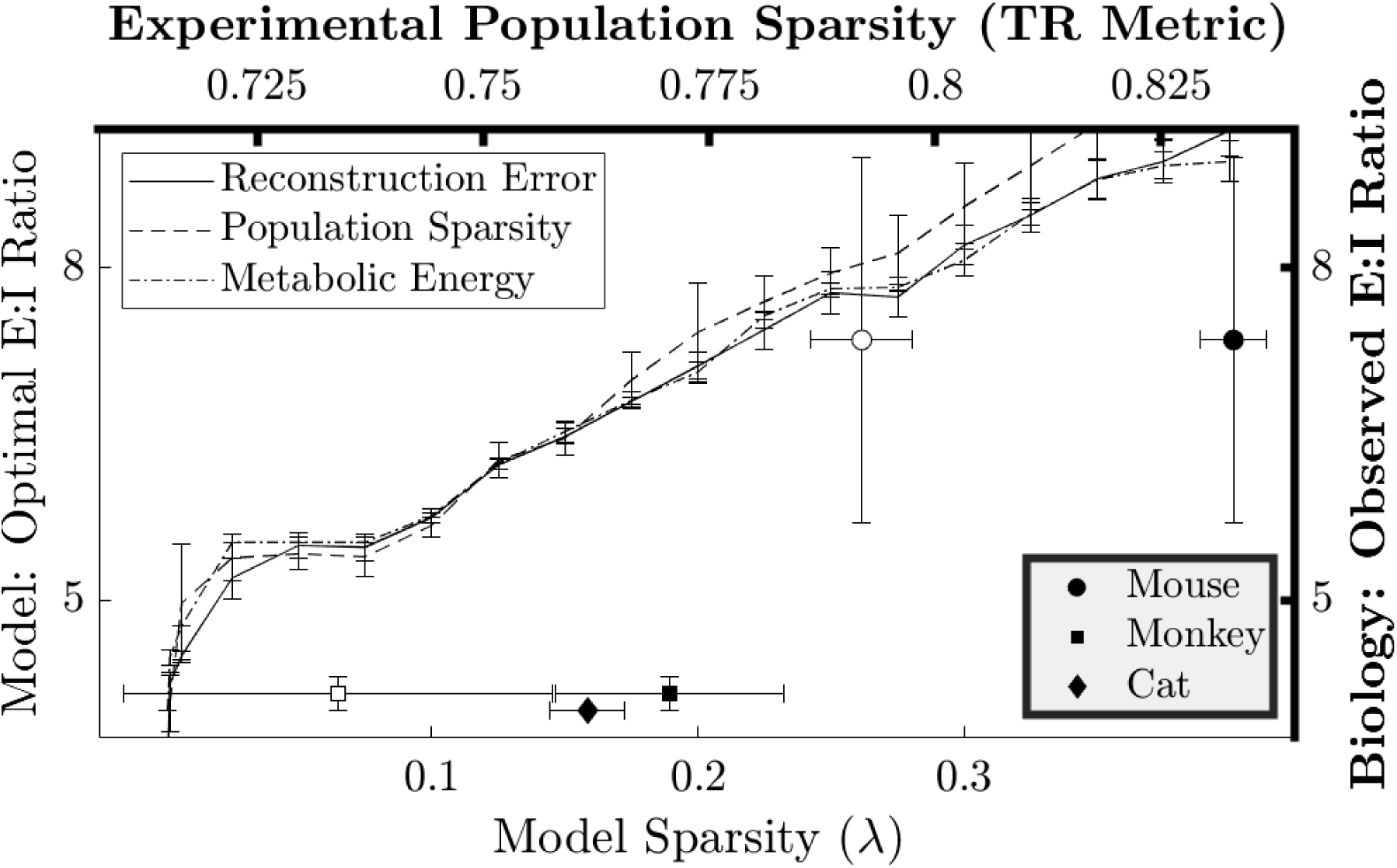
**(Left Y and Bottom X axes)** The Optimal E:I ratio as a function of model sparsity constraints *λ* in computational models according to all three performance measures is captured by the different lines with error bars denoting the standard error for each. A similar (but not identical) trend in the relationship between optimal E:I ratio and model sparsity is revealed for each of the 3 performance measures. **(Right Y and Top X axes)** The Population Sparsity (TR) measure computed for electrophysiology data from experimental studies in mice [3], monkeys [4, 5] and cats [6] is shown as mean (markers) ± standard error(horizontal error bars) w.r.t. observed E:I ratio ranges (vertical error bars) in Biology with unfilled markers representing natural images and black filled markers representing natural movies.

